# Ciliary intrinsic mechanisms regulate dynamic ciliary extracellular vesicle release from sensory neurons

**DOI:** 10.1101/2023.11.01.565151

**Authors:** Juan Wang, Josh Saul, Inna A. Nikonorova, Carlos Nava Cruz, Kaiden M. Power, Ken C. Nguyen, David H. Hall, Maureen M. Barr

## Abstract

Cilia-derived extracellular vesicles (EVs) contain signaling proteins and act in intercellular communication. Polycystin-2 (PKD-2), a transient receptor potential channel, is a conserved ciliary EVs cargo. *Caenorhabditis elegans* serves as a model for studying ciliary EV biogenesis and function. *C. elegans* males release EVs in a mechanically-induced manner and deposit PKD-2-labeled EVs onto the hermaphrodite vulva during mating, suggesting an active release process. Here, we study the dynamics of ciliary EV release using time-lapse imaging and find that cilia can sustain the release of PKD-2-labeled EVs for a two-hour duration. Intriguingly, this extended release doesn’t require neuronal synaptic transmission. Instead, ciliary intrinsic mechanisms regulate PKD-2 ciliary membrane replenishment and dynamic EV release. The ciliary kinesin-3 motor KLP-6 is necessary for both initial and extended ciliary EV release, while the transition zone protein NPHP-4 is required only for sustained EV release. The dihydroceramide desaturase DEGS1/2 ortholog TTM-5 is highly expressed in the EV-releasing sensory neurons, localizes to cilia, and is required for sustained but not initial ciliary EV release, implicating ceramide in ciliary ectocytosis. The study offers a comprehensive portrait of real-time ciliary EV release, and mechanisms supporting cilia as proficient EV release platforms.

## Results and Discussion

Extracellular vesicles (EVs) are submicron membranous structures and key mediators of intercellular communication, shuttling bioactive cargo and signaling molecules to recipient cells^1,2^. Recent research has highlighted roles for cilia-derived EVs in signal transduction, underscoring their importance as bioactive extracellular organelles containing conserved ciliary signaling proteins^3,4^ . Members of the TRP channel Polycystin-2 (PKD-2) family are found in ciliary EVs of the green algae *Chlamydomonas* and the nematode *Caenorhabditis elegans*^5,6^ and in EVs in the mouse embryonic node and isolated from human urine^7,8^.

In *C. elegans*, PKD-2 is exclusively expressed in male-specific EV releasing sensory neurons, which extend their ciliary tips to ciliary pore and directly release EVs into the environment^6,9^. Males release EVs in a mechanically-stimulated manner, regulate EV cargo content in response to mating partners, and deposit PKD-2::GFP-labeled EVs on the vulval cuticle of hermaphrodites during mating^9,10^. Combined, our findings suggest that ciliary EV release is a dynamic process. Herein, we identify mechanisms controlling dynamic EV shedding.

We discovered that cilia are capable of releasing PKD-2::GFP-labeled EVs for a duration of up to two hours. Surprisingly, synaptic transmission is not required for the extended PKD-2 ciliary EV release from sensory neurons. Rather, the ciliary transport machinery involving the kinesin-3 motor KLP-6 and the transition zone protein NPHP-4 are necessary for the replenishment of PKD-2 in the ciliary membrane and this prolonged release of EVs. These findings emphasize the critical role of intrinsic ciliary mechanisms in governing dynamic EV release. We also show that the dihydroceramide desaturase DEGS1/2 ortholog TTM-5 is required for sustained release of PKD-2 ciliary EVs, highlighting the role of ceramide in ciliary ectocytosis. In summary, we visualize real-time ciliary EV release and identify basic principles that govern its dynamic regulation. This work presents a glimpse into how ciliated sensory neurons release EVs in real time in a living animal.

### Extended release of PKD-2::GFP labeled ciliary EVs for two hours

PKD-2-laden ciliary EVs are generated from both the ciliary base and tip, with the latter being directly released into the surrounding environment and mediating inter-animal communication^6,9,11^. *C. elegans* males release PKD-2::GFP-labeled ciliary EVs in response to contact with a coverslip while the animals are mounted on glass slides for microscopic imaging^10^. To gain insight into the dynamics of ciliary EV release, we conducted a series of time-lapse imaging experiments. We found that *C. elegans* males consistently released PKD-2::GFP-labeled ciliary EVs over a span of two hours (Figure 1). We observed environmental EV accumulation at release sites at the head and tail, attributed to the immobilization of animals through anesthesia (Figure 1A). In the head region, EV counts gradually increased, with statistical significance achieved at the two-hour time point (Figure 1B). In the tail region, we observed a more consistent increase in EV numbers, culminating in statistically significant maximum at the one-hour time point (Figure 1B). Further analysis of EV number trajectories of individual animals revealed that, although both the head and tail displayed extended EV release patterns, the tail exhibited a more consistent EV number increase (Figure 1C). We therefore used the more consistent male tail assay for subsequent analysis of EV dynamic release.

**Figure 1.**
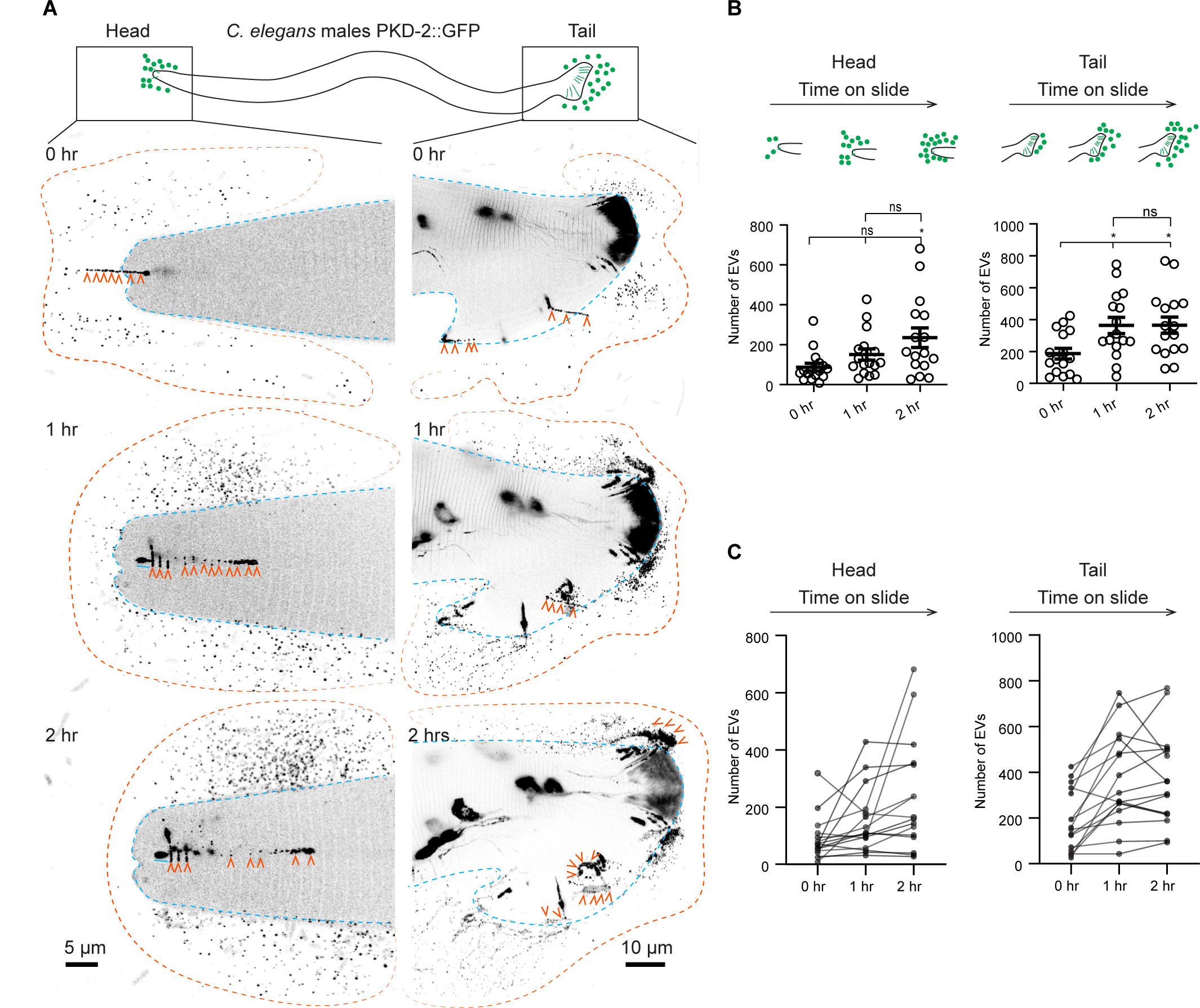
Extended release of PKD-2::GFP-labeled extracellular vesicles (EVs) by *C. elegans* male sensory neurons. (A) Representative images capturing the head and tail regions of *C. elegans* males at 0, 1, and 2 hours following mounting on a glass slide. Blue lines indicate the outline of the male head or tail. Orange lines indicate the outline of the EV clouds released by the head and the tail. Orange arrowheads point to EVs that are released in a string-like pattern. (B) Quantification of EV counts from the head and tail regions at 0, 1, and 2 hours. The scatter plot with lines indicates the mean ± SEM. Each data point represents the total EV count released by an individual *C. elegans* male, either from its head (left) or tail (right). *n* = 16 and the data were scored over 4 days. Statistical analysis was performed using one-way ANOVA with Bonferroni correction. ns denotes not significant (p ≥ 0.05), and * denotes p < 0.05. (C) Individual trajectories depicting the EV release pattern from the head and tail regions of *C. elegans* males. Each trajectory represents EV counts from a single animal. The EV count at new time points includes newly released EVs in addition to previously released EVs that remain visible post-photobleaching. Photobleaching explains the decline in EV counts among certain animals at the 2-hour mark, where newly released EVs are fewer than the photobleached older EVs (photobleached twice). See also Figure S1, Video S1 - S2.

To further characterize the dynamics of EV release during the first hour, we imaged EV release every 10 minutes (Figure S1, Video S1). The number of EVs consistently increased over the course of an hour and reached statistical significance at the 30-minute mark (Figure S1A). Analysis of individual EV trajectories in animals revealed that 5 out of 8 males continuously released EVs over the course of an hour, whereas in 3 out of 8 animals, EV numbers plateaued at the 30-minute mark (Figure S1B). This observation may explain the variability in EV release during the two-hour assay. To visualize real-time EV release, we performed time-lapse imaging for one minute (Video S2, 39 seconds per frame). As the cilium was deflected and moved along the coverslip, the ciliary tip released EVs that appeared as strings and that formed clouds (Figure 1A, Video S2). Our time-lapse imaging confirms that these continuously released EVs originate from the cilia tips. Collectively, these findings demonstrate the capacity of cilia to release EVs for up to two hours from sensory neurons within living animals.

### Synaptic transmission is not required for dynamic PKD-2 ciliary EV release from sensory neurons

Sensory neurons release EVs from cilia when mechanically stimulated and during mating^10^. Therefore, we asked whether synaptic transmission and communication from other neurons was necessary for the dynamic PKD-2 ciliary EV release. We examined mutants defective in docking synaptic vesicles (*unc-13*) and in dense core vesicle exocytosis (*unc-31*)^12–14^ (Figure 2). Neither *unc-13* nor *unc-31* mutant males displayed deficiencies in PKD-2 ciliary EV release (see Figure 2A-C). At the initial time point (0 hours), both *unc-13* and *unc-31* mutants exhibited PKD-2 EV release at wild-type levels. The cumulative EV release over one hour was comparable between *unc-13*, *unc-31*, and wild-type males (Figure 2). This data indicates that synaptic vesicle and dense core vesicle exocytosis are not required for PKD-2 initial or extended ciliary EV release.

**Figure 2.**
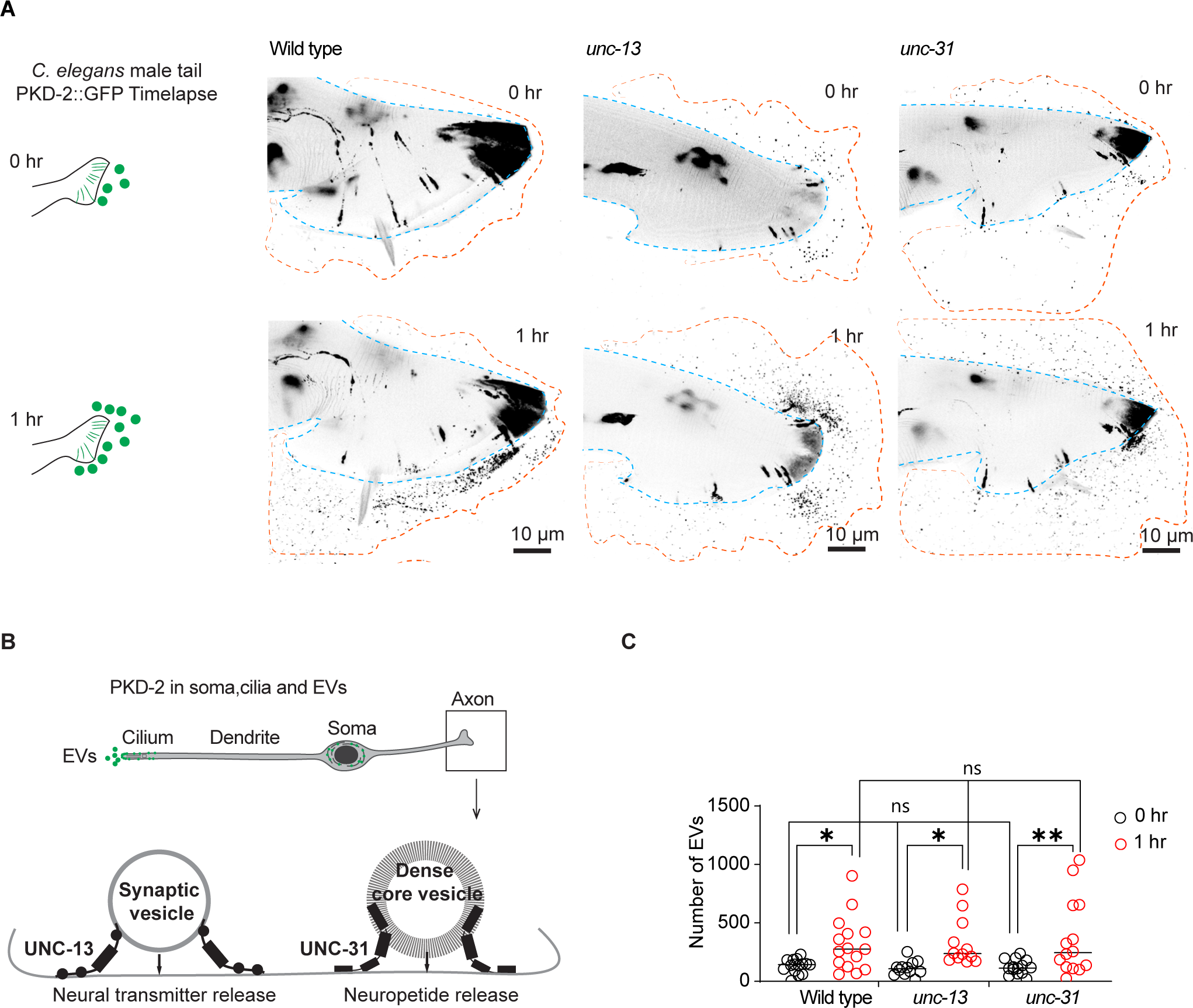
Neuronal transmission-independent release of PKD-2::GFP-labeled ciliary EVs. (A) Representative images showing EV release from the tail at the initial imaging (0 hr) and the second imaging (1 hr) while the animals were mounted on slides, in wild-type, *unc-13*, and *unc-31* males. (B) Schematic diagram depicts the functional roles of UNC-13 and UNC-31 in synaptic vesicle- and dense core vesicle-mediated neuronal transmission in the axon. (C) Quantification of EV release from the tail at the initial imaging (0 hr) and second imaging (1 hr) in wild-type, *unc-13*, and *unc-31* males. The scatter plot with lines indicates the mean ± SEM. Each data point represents the total EV count released by an individual *C. elegans* male. 12-15 animals were imaged for each genotype. Statistical analysis was performed by two-way ANOVA with Bonferroni correction. ns denotes not significant, * denotes p < 0.05, and ** denotes p < 0.01.

### Continuous EV release relies on the membrane replenishment of PKD-2 at the ciliary tip

The ciliary kinesin-3 motor KLP-6 (<u>k</u>inesin like <u>p</u>rotein <u>6</u>) is essential for PKD-2 environmental EV release at the ciliary tip. In *klp-6* mutant animals, PKD-2 is not released in environmental EVs, leading to excessive shedding of EVs at the ciliary base, into the glial lumen surrounding the ciliary base^6^ (Figure S2A-D). To explore the potential contribution of the ciliary base EV reservoir to the sustained release of PKD-2 EVs, we conducted a time-lapse analysis of PKD-2 EV release in *klp-6* mutant males (Figure 3). *klp-6* mutant animals exhibited a defect in environmental EV release at both the initial and the one-hour time point (Figure 3A, B, D).

**Figure 3.**
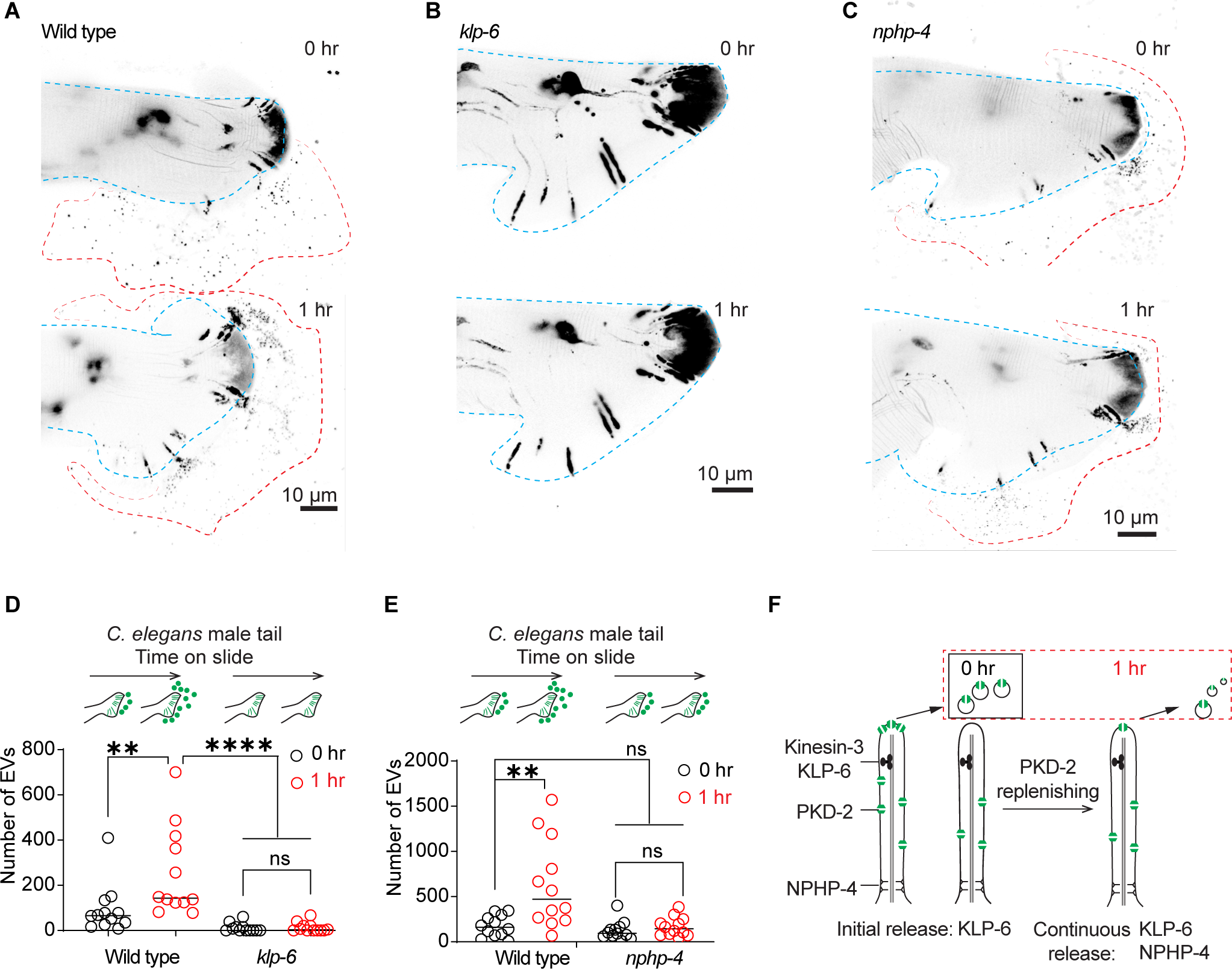
Continued PKD-2 ciliary EV release relies on effective replenishment of PKD-2 at the ciliary tip. (A-C) Representative images of PKD-2::GFP ciliary EV release in male tail of wild-type, *klp-6*, and *nphp-4* males. (D) Quantification of EV counts from wild-type and *klp-6* mutant males in the tail regions at 0 and 1 hour. The scatter plot with lines indicates the mean ± SEM. Each data point represents the total EV count released by an individual *C. elegans* male. (E) Quantification of EV counts from wild-type and *nphp-4* mutant males in the tail regions at 0 and 1 hour. The scatter plot with lines indicates the mean ± SEM. Each data point represents the total EV count released by an individual *C. elegans* male. For D-E, statistical analysis was performed by two-way ANOVA with Bonferroni correction. n = 12-15 animals per genotype. ns denotes not significant, * denotes *p* < 0.05, and ** denotes *p* < 0.01., **** p<0.0001. (F) Model of PKD-2 ciliary EV dynamics. NPHP-4 regulates the release of PKD-2::GFP ciliary EVs by replenishing PKD-2::GFP in the ciliary membrane. Meanwhile, KLP-6 controls ciliary PKD-2 EV release by concentrating PKD-2 at the ciliary tip. The sustained release of PKD-2 ciliary EVs over an hour necessitates the replenishment of PKD-2 in both the ciliary membrane and tip. See also Figure S2.

These data show that ciliary base EVs are not released into the environment during the prolonged one-hour time-lapse assay, and that KLP-6 kinesin motor-facilitated enrichment of EV cargos at the ciliary tip is necessary for the dynamic release of PKD-2 ciliary EVs.

To test the hypothesis that continuous EV release originates from the ciliary tip, we reasoned that a mutant defective in PKD-2 ciliary membrane replenishment would also be defective in sustained EV release (model shown in Figure 3F). In cultured mammalian cells, the KLP-6 homolog KIF13B physically interacts with the transition zone protein NPHP4 (<u>n</u>e<u>ph</u>rono<u>p</u>hthisis gene <u>4</u>) to regulate ciliary membrane composition and Sonic hedgehog signaling^15^. Therefore, we determined whether NPHP-4 was important for PKD-2 ciliary EV release. We previously found that the *nphp-4* mutant does not grossly affect PKD-2 ciliary localization^16^. At the initial time point, the *nphp-4* mutant did not show defects in PKD-2::GFP ciliary EV release. However, at the one-hour time point, *nphp-4* mutant males failed to release additional EVs (Figure 3 C, F). Using transmission electron microscopy (TEM), we found that *nphp-4* mutants displayed excessive EV accumulation surrounding the ciliary base (Figure S2E-F). The *nphp-4* ciliary base EV accumulation phenotype resembles the *klp-6* mutant phenotype (Figure S2), consistent with the two genes acting in a common process. The finding that the *nphp-4* mutant is required for sustained PKD-2 ciliary EV release, but not for initial EV release, suggests that the prolonged PKD-2 EV release requires efficient ciliary replenishment of PKD-2. Like their mammalian homologs, NPHP-4 might interact with KLP-6 to facilitate the efficient import of ciliary membrane proteins. The transition zone plays important roles in regulating ciliary protein and lipid content^17^, the latter raising the question of the potential function of lipids in ciliary EV biogenesis.

### Dihydroceramide desaturase TTM-5 is required for the extended PKD-2 ciliary EV release

The sustained release of ciliary EVs requires an abundant supply of membrane lipids. As per our estimation, the membrane of approximately 60 EVs equates to the ciliary membrane (Figure S3A). Ceramide is a pivotal membrane lipid essential for EV formation^2,18^. Ceramide is also an important lipid for cilia formation, ciliary length and ciliary signaling^19^. TTM-5 (Toxin-regulated Targets of MAPK 5) is an ortholog of human DEGS1/DEGS2 (delta 4-desaturase, sphingolipid) that plays a role in de novo ceramide synthesis^20–22^ (Figures 4A). *ttm-5* shows high expression in IL2 ciliated sensory neurons^23^, which are the sex-shared EV-releasing neurons (Figure S3B) and express ciliary EV biogenesis genes such as *klp-6* and *cil-7* ^24–26^.

**Figure 4.**
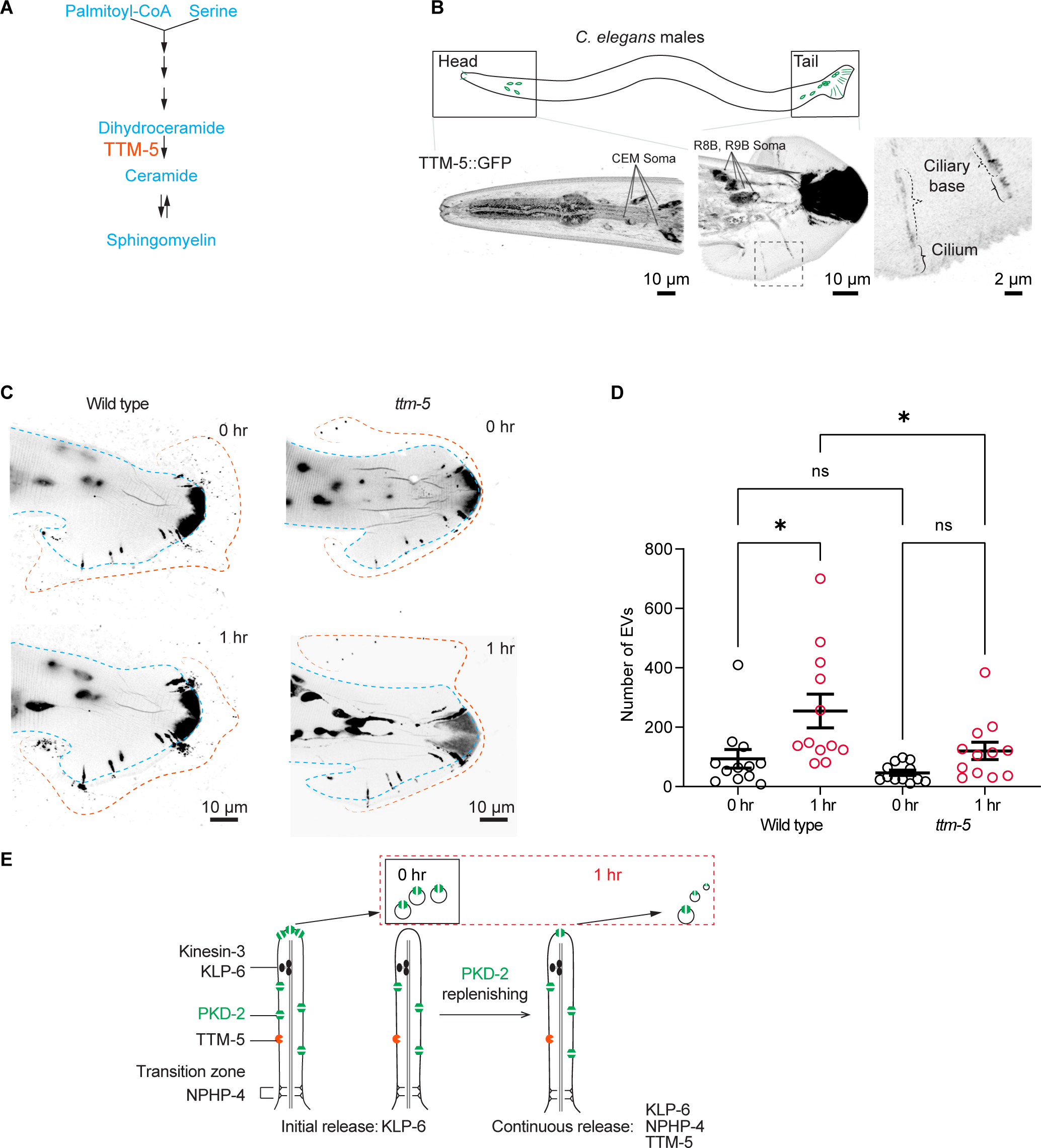
TTM-5 facilitates the dynamic release of PKD-2 ciliary EVs. (A) Schematic cartoon of TTM-5 action site. TTM-5 is the ortholog of DEGS1/2 (delta-4-desaturase, sphingolipid 1/2), an enzyme that converts dihydroceramides to ceramides. The unsaturated fatty acid chain creates a kink in the chain, making the cone shape of mature ceramides to support the high curvature membrane of cilia and EV. (B) A CRISPR reporter of TTM-5::GFP is expressed in male-specific EV-releasing neurons, including four cephalic male (CEM) neurons and 17 neurons in the tail (soma of R8B and R9B are shown, for the full tail images, see Figure S3C). In the tail EV-releasing neurons, TTM-5 primarily localizes to the soma, ciliary base, and cilium and is not detectable in EVs. (C) Representative images of PKD-2::GFP ciliary EV release in male tail sensory neurons at 0 and 1 hour in wild-type and *ttm-5* mutant males. Blue lines indicate the outline of the male tail. Orange lines indicate the outline of the EV clouds released by the tail. (D) Quantification of EV counts at 0 and 1 hour of wild-type and *ttm-5* mutant males. The scatter plot with lines indicates the mean ± SEM. Each data point represents the total EV count released by an individual *C. elegans* male. n = 12. Statistical analysis was performed using one-way ANOVA with Bonferroni correction. ns denotes not significant (p ≥ 0.05), and * denotes p < 0.05. See also Figure S3-4.

To test whether TTM-5 might function in the PKD-2 EV releasing neurons, we edited the *ttm-5* genomic location using a CRISPR/Cas9 based methodology to generate an endogenous reporter by fusing *ttm-5* to the GFP encoding sequence. TTM-5::GFP is primarily expressed in the EV-releasing neurons as demonstrated by co-expression of TTM-5::GFP with LOV-1::mSc^27^, as well as the single reporter TTM-5::GFP. In the male head, TTM-5 is expressed in IL2 and CEM neurons, along with several amphid neurons. In the male tail, TTM-5 is coexpressed with LOV-1 in HOB and RnB neurons (Figures 4B and Figure S3C). TTM-5::GFP localizes to the cilia; however, TTM-5 is not discernibly present in ciliary EVs (Figures 4B and Figure S3C). This is consistent with our prior EVome profiling data from which TTM-5 is absent^28^.

TTM-5 expression in the EV-releasing neurons and localization in cilia suggest a role in ciliary EV biogenesis. In the *ttm-5(tm6585)* loss-of-function mutant (Figure S4), the localization of PKD-2::GFP appears largely normal (Figure 4C). We next examined EV release. In the *ttm-5* mutant, initial EV release was comparable to wild type. However, the *ttm-5* mutant displayed an inability to release additional EVs in the one-hour assay (see Figures 4C-D). In summary, our findings indicate that the enzyme TTM-5, associated with the ceramide metabolism pathway, is enriched in the EV-releasing neurons, localizes to the ciliary membrane, and is vital for the sustained release of PKD-2 ciliary EVs.

*C. elegans* males release ciliary EVs in response to mechanical stimuli and in response to mating partners^9,10^. Here we comprehensively characterize real-time ciliary EV release and identify components that govern its dynamic regulation. Our study offers novel insights into the dynamic regulation of EV release from cilia of sensory neurons in *C. elegans*. The prolonged release of PKD-2::GFP-labeled ciliary EVs for up to two hours underscores the robust capability of cilia to produce EVs. This work opens new avenues for exploring the regulators of ciliary EV biogenesis, cargo sorting, and signaling.

Synaptic transmission is not required for dynamic PKD-2 ciliary EV release. This finding is consistent with a previous report showing that touch-induced calcium responses in these ray RnB neurons do not rely on synaptic transmission^29^. This ciliary EV shedding, independent of synaptic transmission, aligns with the finding that ciliary EVs carrying polycystin-2 are released from single-celled *Chlamydomonas*^5^. Our results are consistent with a ciliary intrinsic mechanism mediating the dynamic release of PKD-2 ciliary EVs.

Both UNC-13 and UNC-31 are conserved bridge molecules between fusing vesicles and target membranes. Neither *unc-13* nor *unc-31* is required for PKD-2 ciliary EV release. However, we do not rule out that *unc-13* and *unc-31* regulate MVB-mediated exosome biogenesis in *C. elegans*. The *Drosophila* UNC-13 homolog, stac, localizes to the multivesicular body (MVB) and is required for MVB-mediated EV release and tracheal cell fusion in embryos^30^. The human ortholog of UNC-13, Munc13-4, is required for MVB maturation and exosome release in cancer cells^31^. The UNC-31 ortholog, CAPS1, overexpression promotes exosome-regulated cancer cell migration^32^.

The requirement for KLP-6 in extended EV shedding is in line with previous findings indicating KLP-6’s essential role in EV shedding from the ciliary tip^6,9^. This observation supports the hypothesis that prolonged PKD-2 ciliary EV release originates from the ciliary tip rather than the base EVs. Within primary cilia, NPHP4 interacts with KIF13B, a KLP-6 homolog, to regulate ciliary membrane composition and Hedgehog signaling^15^. Intriguingly, NPHP4 also plays a role in whole cilia shedding, where cilia break off at the transition zone in *Paramecium*^33^. In *Chlamydomonas,* NPHP4 and other transition zone components influence the proteome of ciliary ectosomes^34^. We propose a cilia-intrinsic regulatory mechanism whereby NPHP-4 operates through KLP-6 to facilitate PKD-2 ciliary membrane replenishment, thus supporting effective PKD-2 ciliary EV release (Figure 4E). Furthermore, our findings highlight a role for NPHP-4 as a conserved regulator of ciliary EV biogenesis. Mutations in transition zone components like NPHP4 cause human ciliopathies^35^, suggesting that EV defects may contribute to ciliopathies such as nephronophthisis and polycystic kidney disease^4^.

TTM-5 is a dihydroceramide desaturase (DEGS1/2) ortholog, which acts in a ceramide biosynthesis pathway to convert dihydroceramide to ceramide^22^. Ceramide is a bioactive signaling lipid that plays roles in ciliogenesis, ciliary TGF-β receptor/sonic hedgehog signaling, and exosome biogenesis^2,19^. With respect to the latter, the cone-shaped structure of ceramide induces negative membrane curvature that leads to invagination into the endosome membrane and formation of inner luminal vesicles of the MVB – a prerequisite for exosome biogenesis. Here, we show that TTM-5 is required for the sustained, but not initial, release of ciliary-derived EVs. We propose that, similar to exosome biogenesis, the cone shape of ceramide induces membrane curvature and ectocytosis from the ciliary tip.

The high expression of *ttm-5* specifically in EV releasing neurons and the ciliary localization of TTM-5 suggests that EV releasing neurons optimize a membrane lipid synthesis pathway for sustained ciliary EV release and constant renewal of the ciliary membrane. Vertebrate photoreceptors generate an enormous amount of light sensitive membrane daily and are capable of producing vast quantities of ectosomes^36,37^. The ability to form ectosome is blocked by peripherin-2 to allow outer segment formation^38^. Like photoreceptors, the *C. elegans* EV releasing neurons produce copious amounts of ciliary EVs. The DEGS1/2 TTM-5-mediated lipid synthesis pathway may be important for generating the ciliary membrane required to support sustained release of EVs. Sphingolipid metabolism enzymes play a conserved role in cilium formation in *Chlamydomonas*, *C. elegans*, and vertebrate cells, linking compromised sphingolipid production with ciliopathies^39^.

Our previous super-resolution imaging studies revealed that cilia release EVs from two sites – the cilia base and tip^9^. Here we developed a powerful one-hour imaging approach that provides spatiotemporal information on ciliary EV shedding in real-time. This extended imaging strategy revealed that different mechanisms control initial versus prolonged EV release. Not surprisingly, kinesin-3 KLP-6 is required for both initial and prolonged EV release. In contrast, the transition zone component NPHP-4 and the dihydroceramide desaturase TTM-5 are important for prolonged but not initial EV release. These results suggest that the transition zone – the ciliary gate keeper – and the sphingolipid ceramide play important roles in maintaining steady-state ciliary ectocytosis. Our *C. elegans* system enables discovery of conserved mechanisms regulating ciliary membrane renewal and ciliary ectocytosis, which provides much needed in vivo insight to EV biogenesis and signaling.

## Acknowledgments

This work was supported by National Institutes of Health (NIH) DK059418, DK116606 and NS120745 (M.M.B). We thank Gloria Androwski for excellent technical assistance, Barr lab mates and the Rutgers *C. elegans* community for feedback and constructive criticism throughout this project. We also thank WormBase, Japan National Bioresource Project for the nematode and Caenorhabditis Genetics Center (CGC) for resources information and strains. The CGC is supported by the National Institutes of Health - Office of Research Infrastructure Programs (P40OD010440).

## Author contributions

Conceptualization, JW; Methodology, JW, JS, IAN, KCN, DHH, MMB; Investigation, JW, JS, IAN, CNC, KMP; Visualization, JW, JS, IAN, CNC, KMP; Funding acquisition, DHH and MMB; Project administration, JW, IAN, and MMB; Supervision, JW, DHH, MMB; Writing – original draft, JW; Writing – review & editing, JW, JS, IAN, and MMB.

## STAR★METHODS

### Key Resources Table

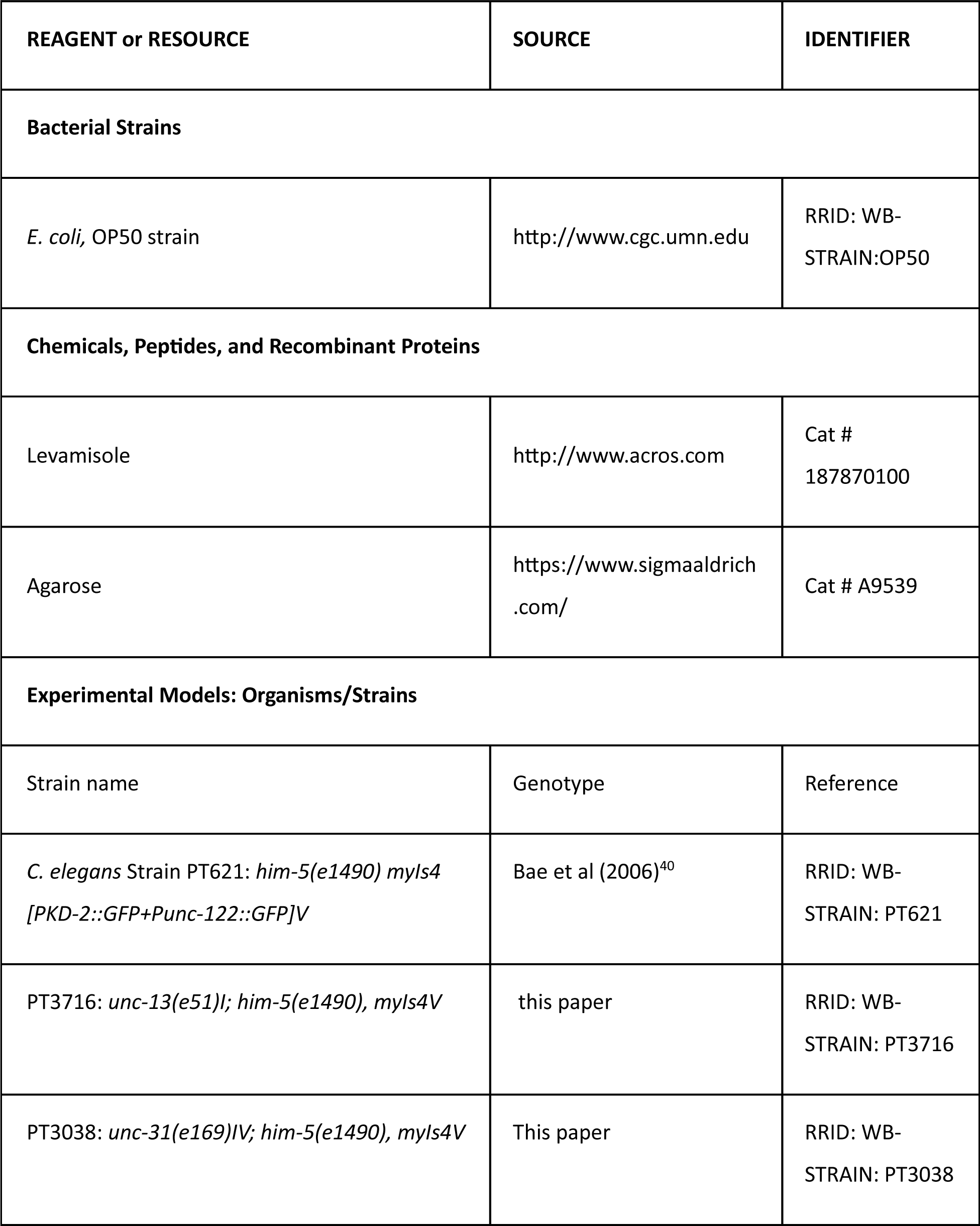
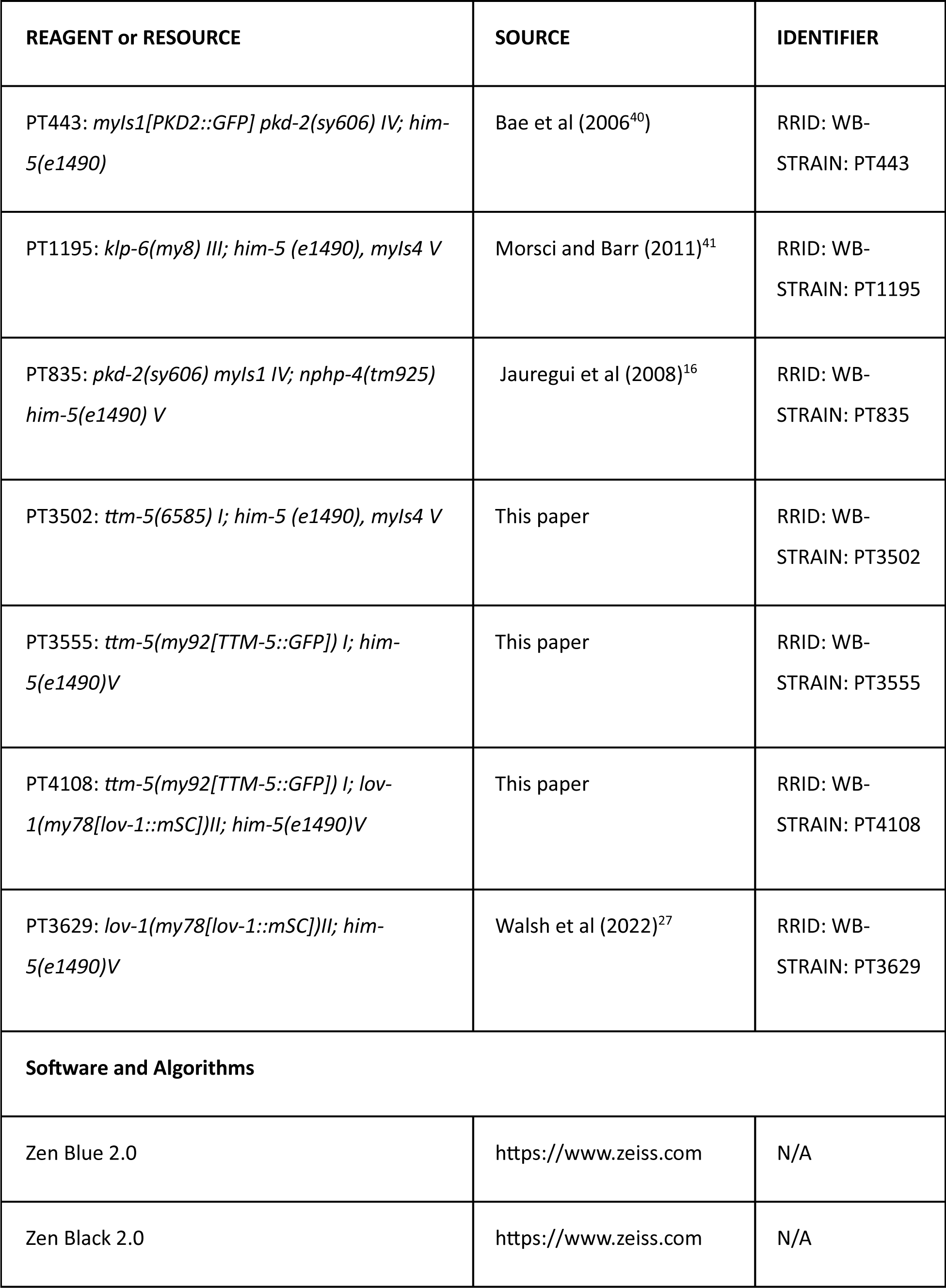
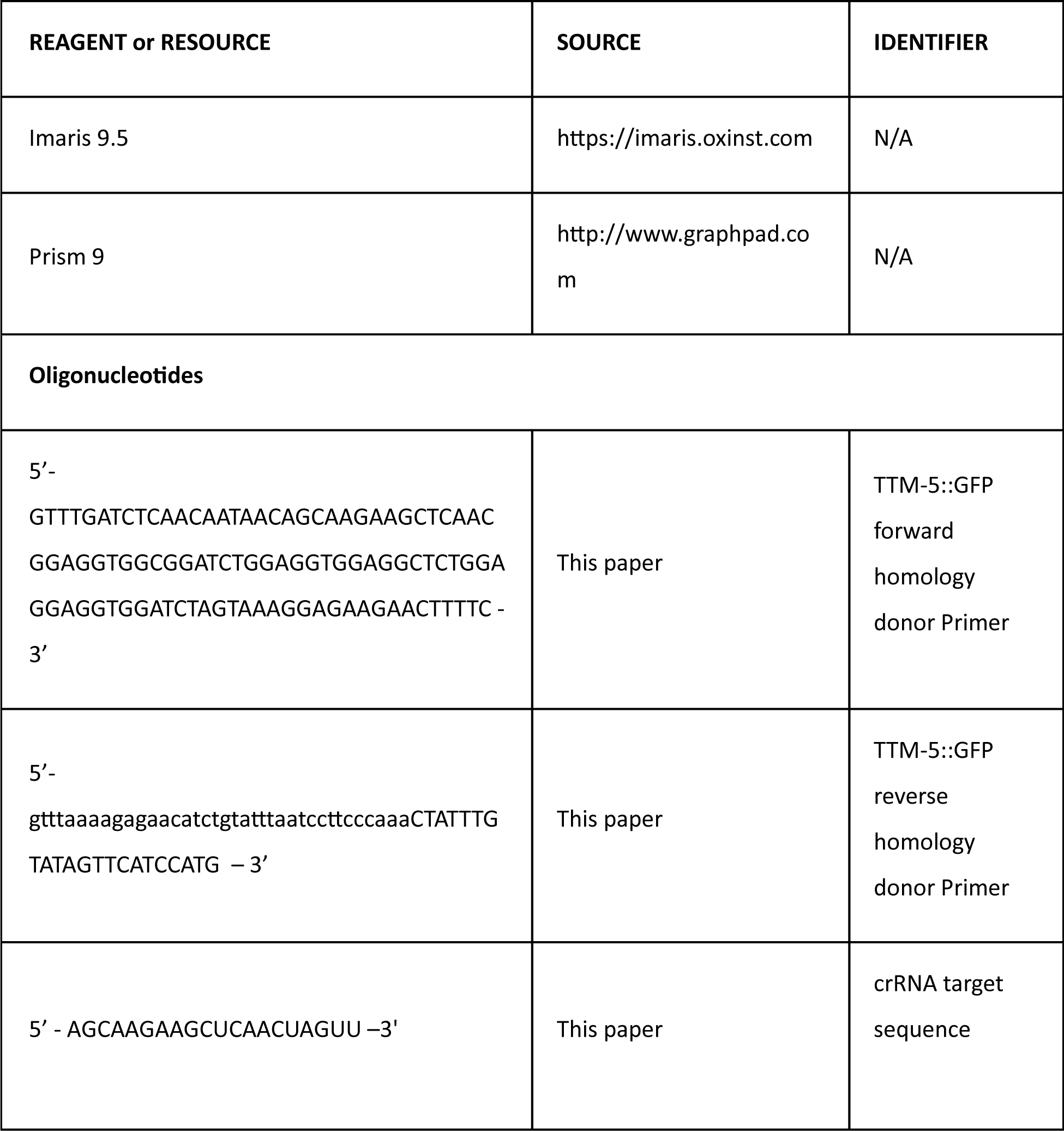

#### Lead contact

Further information and requests for resources and reagents should be directed to and will be fulfilled by the lead contacts Juan Wang and Maureen Barr.

#### Materials availability

Plasmids and transgenic *C. elegans* strains are available upon request.

#### Data and code availability

Microscopy data reported in this paper will be shared by the lead contact upon request.

Any additional information required to reanalyze the data reported in this paper is available from the lead contact upon request.

### Experimental model and subject details

*C. elegans* culture and genetics were performed as described^42^. *C. elegans* strains were maintained at 20°C on nematode growth medium (NGM) plates seeded with *Escherichia coli* (OP50 strain) as a food source.

Strain identifiers and genotypes are cataloged in the Key Resources Table.

## Method details

### Airyscan super-resolution microscopy

Super-resolution imaging was performed on the Zeiss LSM880 confocal system equipped with Airyscan Super-Resolution Detector with 7 Single Photon Lasers (405, 458.488, 514, 561, 594, 633nm), Axio Observer 7 Motorized Inverted Microscope, Motorized X-Y Stage with Z-Piezo, T-PMT.

### Time-lapse imaging and PKD-2::GFP EVs quantification

The day before imaging, 20-30 L4 larval males were picked to NGM plates freshly seeded with *E. coli* OP50 to attain a synchronized population. On the day of imaging, now-adult male *C. elegans* were picked from the NGM plates to a coverslip containing 1 μL drops of 10 mM levamisole in water and mounted on a 10% agarose pad on a slide, and imaged in 1-minute, 10-minute or one-hour intervals. All images were taken with Airyscan super-resolution on an LSM 880 confocal microscope (Zeiss). Images were saved and processed using Zen Black software (Zeiss) to generate maximum intensity projections and Zen Blue software (Zeiss) image analysis wizard to quantify the number of EVs. Images were processed and prepared for publication using Imaris, FIJI software, Adobe Illustrator and Adobe Photoshop. Maximum intensity projects of Z-stack images were inverted and adjusted to grayscale in Adobe Photoshop.

### Transmission Electron Microscopy (TEM)

For method details, refer to Jauregui et al (2008)^16^. Briefly, worms were fixed in 3% glutaraldehyde in cacodylate buffer on ice. Then, heads were cut off, moved to fresh fixative, and held in fix overnight at 4°C. Animals were rinsed in buffer and stained with 1% osmium tetroxide in cacodylate buffer for 1 hr at 4°C. After embedding in small groups in agarose, the specimens were en bloc stained with 1% uranyl acetate in sodium acetate buffer, dehydrated, and embedded in Embed812 resin according to the general procedures described by Hall (1995)^43^. Thin sections were collected on a diamond knife and post-stained before being viewed on a CM10 electron microscope.

### Quantification and statistical analysis

The Prism software package (GraphPad Software 8) was used to carry out statistical analyses. Information about statistical tests, p values and n numbers are provided in the respective figures and figure legends.

### CRISPR/Cas9-mediatad editing

To generate the TTM-5::GFP CRISPR endogenous CRISPR tags, we followed the Mello Lab protocol and used partially single-stranded dsDNA donors^44^ with 35-bp of flanking homology^45^ that included a short flexible linker sequence between the 3’ end of *ttm-5* and the start of GFP. The guide sequence was designed using CRISPOR^46^ [http://crispor.tefor.net/].

### Supplemental Video and Data Title and Legend

**Video S1, Timelapse capture of dynamic EV release at 10-minute intervals per frame, related to** **Figure 1**.

The video presents time-lapse imaging captured at 10-minute intervals over a one-hour period, demonstrating the continuous release of EVs throughout that duration. The first frame displays a cartoon of a *C. elegans* male; the rectangle indicates the imaging area for the micrograph shown below. The micrograph is an overlap of bright field and a fluorescence image of the male tail, demonstrating that PKD-2::GFP is enriched at the ciliary tip and is released from the ray pore structure into the environment outside the worm. The image is adapted from Figure 1 in Wang et al (2020)^10^.

**Video S2, related to** **Figure 1. Timelapse capture of dynamic EV release at 39 second intervals per frame.**

The video presents time-lapse imaging captured at 39-second intervals over five minutes demonstrating the continuous release of EVs from the ciliary tip. The left panel displays a cartoon of a *C. elegans* male; the rectangle indicates the imaging area for the micrograph shown on the right. The micrograph is an overlap of bright field and a fluorescence image of the male tail, demonstrating that PKD-2::GFP is enriched at the ciliary tip and is released from the ray pore structure into the environment outside the worm. The image is adapted from Figure 1 in Wang et al (2020)^10^.

---

**Figure S1.**
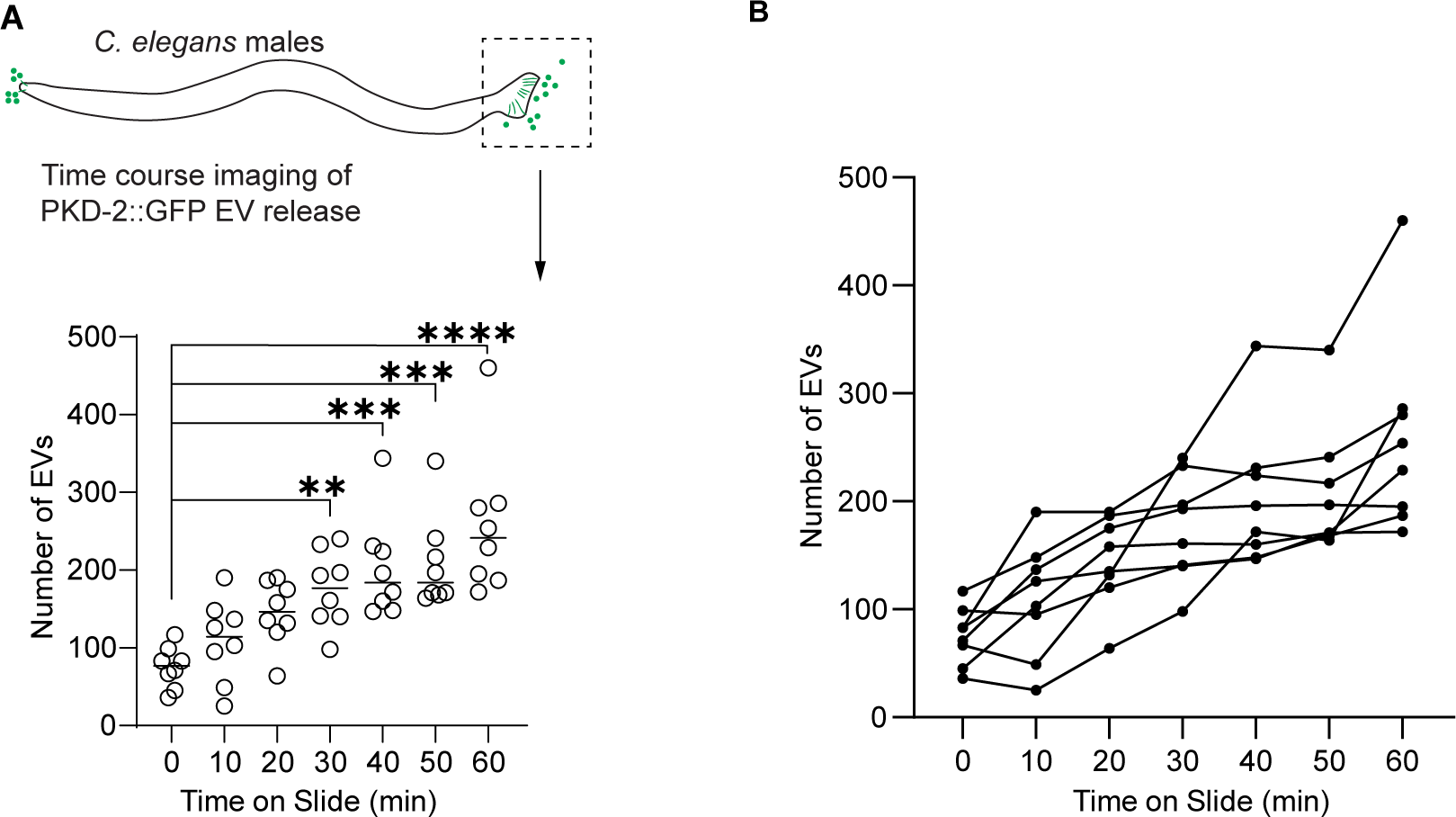
PKD-2::GFP ciliary EV release dynamics, related to Figure 1. (A) Quantification of EV numbers during time-lapse imaging of an hour. Each circle represents total number of EVs released by one animal from the tail at the time point. n= 8 animals. Statistical analysis was performed using one-way ANOVA with Bonferroni correction. ** denotes p < 0.01 and **** denotes p < 0.0001. (B) Individual trajectories depicting the EV release pattern from the male tail at various time points. Each trajectory represents EV counts from a single animal. The EV count at new time points includes newly released EVs in addition to previously released EVs that remain visible post-photobleaching. Photobleaching may explain the decline in EV counts among certain animals where newly released EVs are fewer than previously released EVs, which would have been photobleached multiple times.

**Figure S2.**
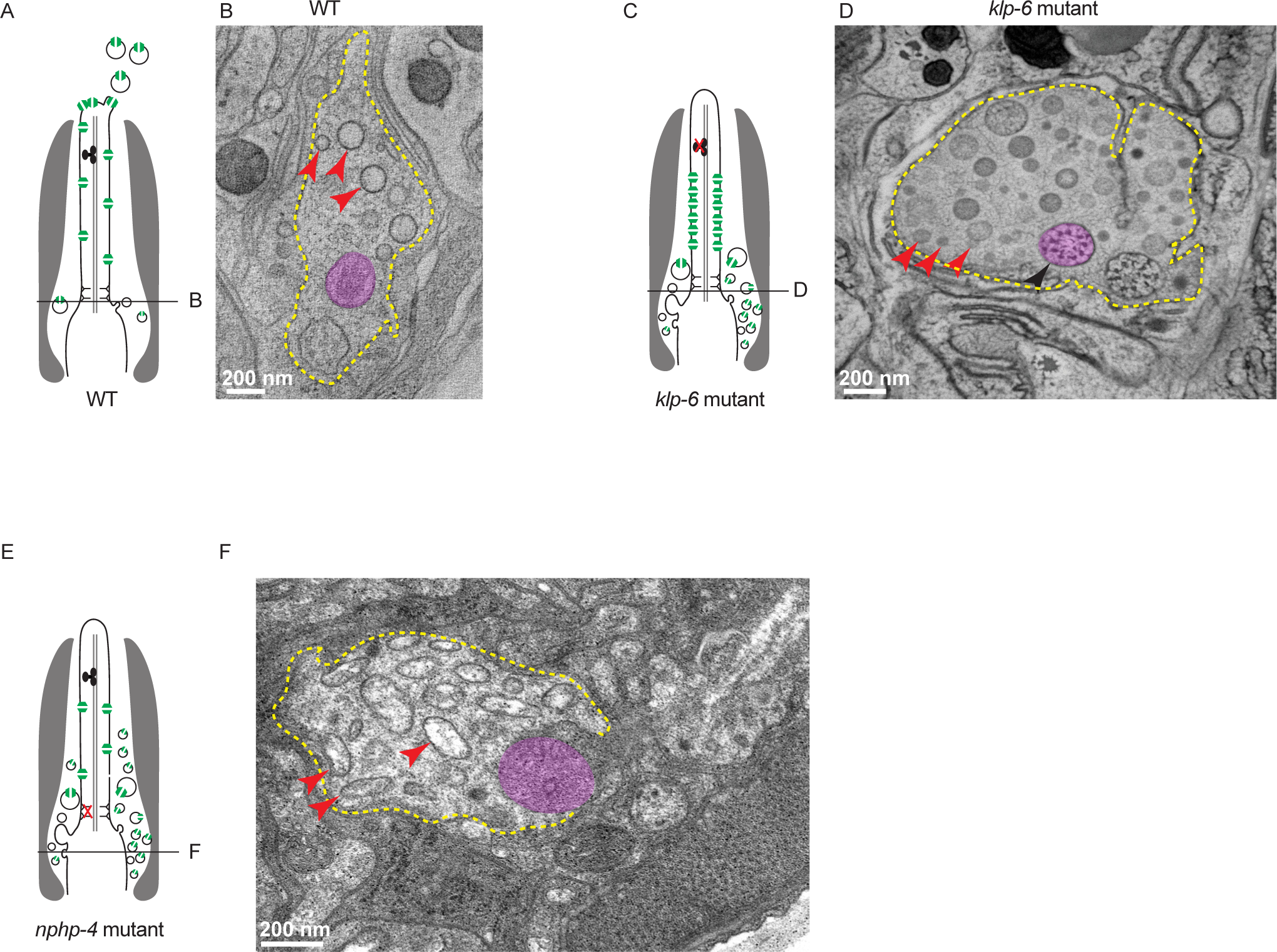
*nphp-4* mutant has excessive internal EVs, related to **Figure 3**. (A, C, E) Schematic representation of a cephalic male (CEM) cilium, illustrating its relationship to the ciliary tip and base EVs in wild type, *klp-6* and *nphp-4* mutants respectively. The environmentally-released EVs originate from the ciliary tip, while the ciliary base EVs are located in the lumen formed by the surrounding glia (depicted in gray). Green circle with a channel denotes PKD-2. Black lines indicate the approximate position of the cross-section transmission electron microscopy (TEM) image shown in corresponding panels. Modified from [S1-2]. (B, D) TEM image of a cross section through a ciliary region in wild-type and *klp-6* mutant respectively . The *klp-6* mutant accumulates excess EVs in the lumen (reproduced from [S1]). (F) TEM image of *nhph-4* mutant ciliary region. In the *nphp-4* mutant, more EVs accumulates in the lumen surrounding the ciliary base. TEM image is from the same TEM series shown in Figure 4i [S3]. In all TEM images, the red arrowheads indicate EVs. The CEM cilium is highlighted in magenta. The glial lumen encircling the CEM cilium is outlined by a yellow dashed line. Scale bars represent 200 nm.

**Figure S3.**
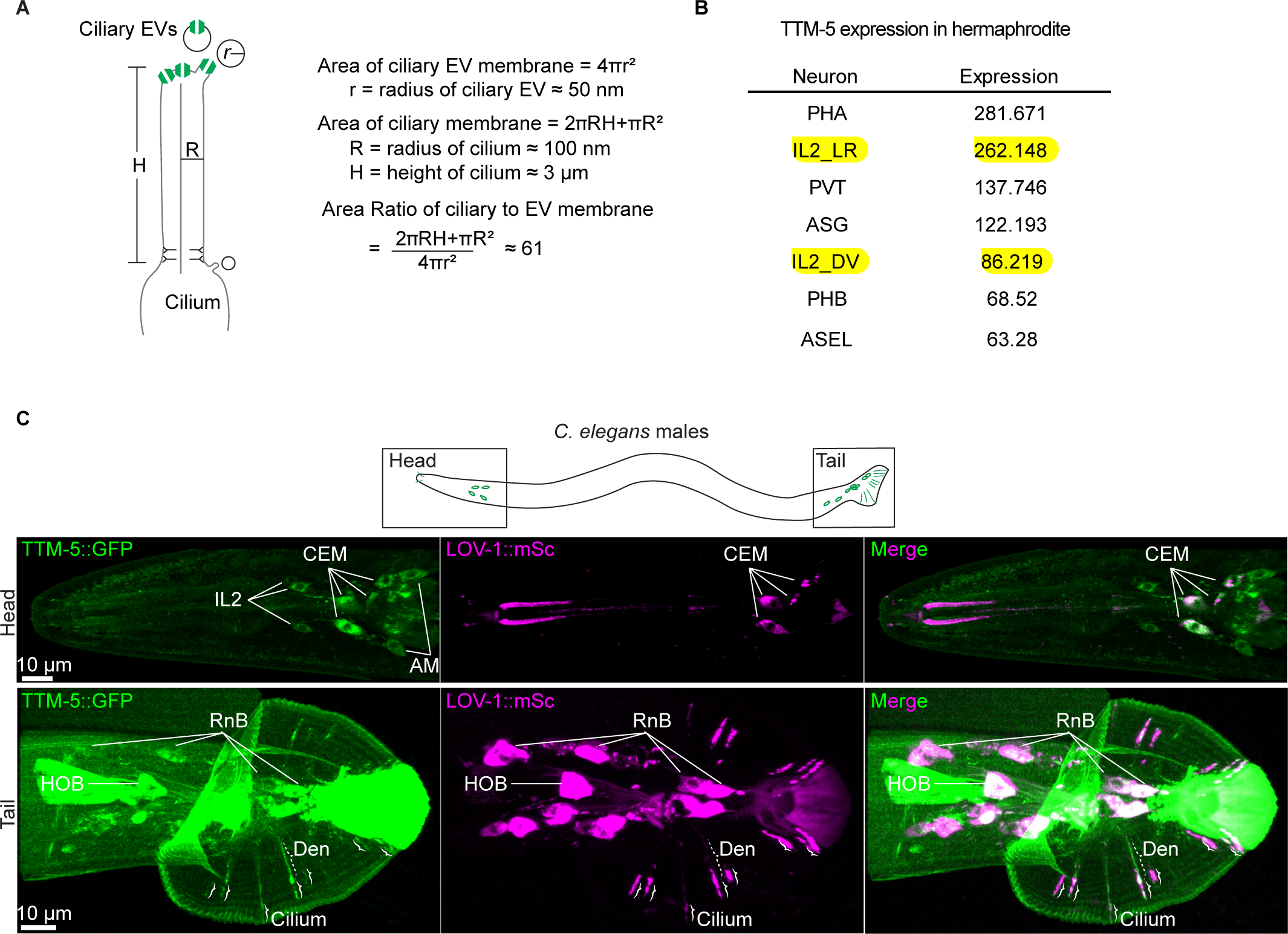
Estimation of ciliary EV membrane requirements and analysis of *ttm-5* expression, related to **Figure 4**. (A) A calculation of the ratio between ciliary and EV membrane surface areas. (B) Single cell RNAseq data from CENGEN APP (CeNGENApp) listed IL2 neurons, the hermaphrodite EV releasing neurons as top expressing neurons of *ttm-5* in the hermaphrodite *C. elegans* [S4]. (C) Images of *C. elegans* males showing the co-expression of TTM-5::GFP (green) and LOV-1::mSc (magenta) in male specific EV releasing neurons. Top panel, schematic of *C. elegans* male-specific EV releasing neurons. The black rectangles indicate the imaged areas in the head and the tail. Lower panel, In the head, TTM-5::GFP is expressed in the inner labial type 2 (IL2) neurons, the cephalic male (CEM) neurons, and some unidentified amphid (AM) neurons. In the tail, TTM-5 is expressed in the male-specific EV releasing neurons, the HOB, and ray type B (RnBs) neurons. TTM-5::GFP localizes to the cilia in all EVNs but is not visibly detected in EVs. Dashed bracket indicate the dendrite (Den), and solid brackets indicate cilia.

**Figure S4.**
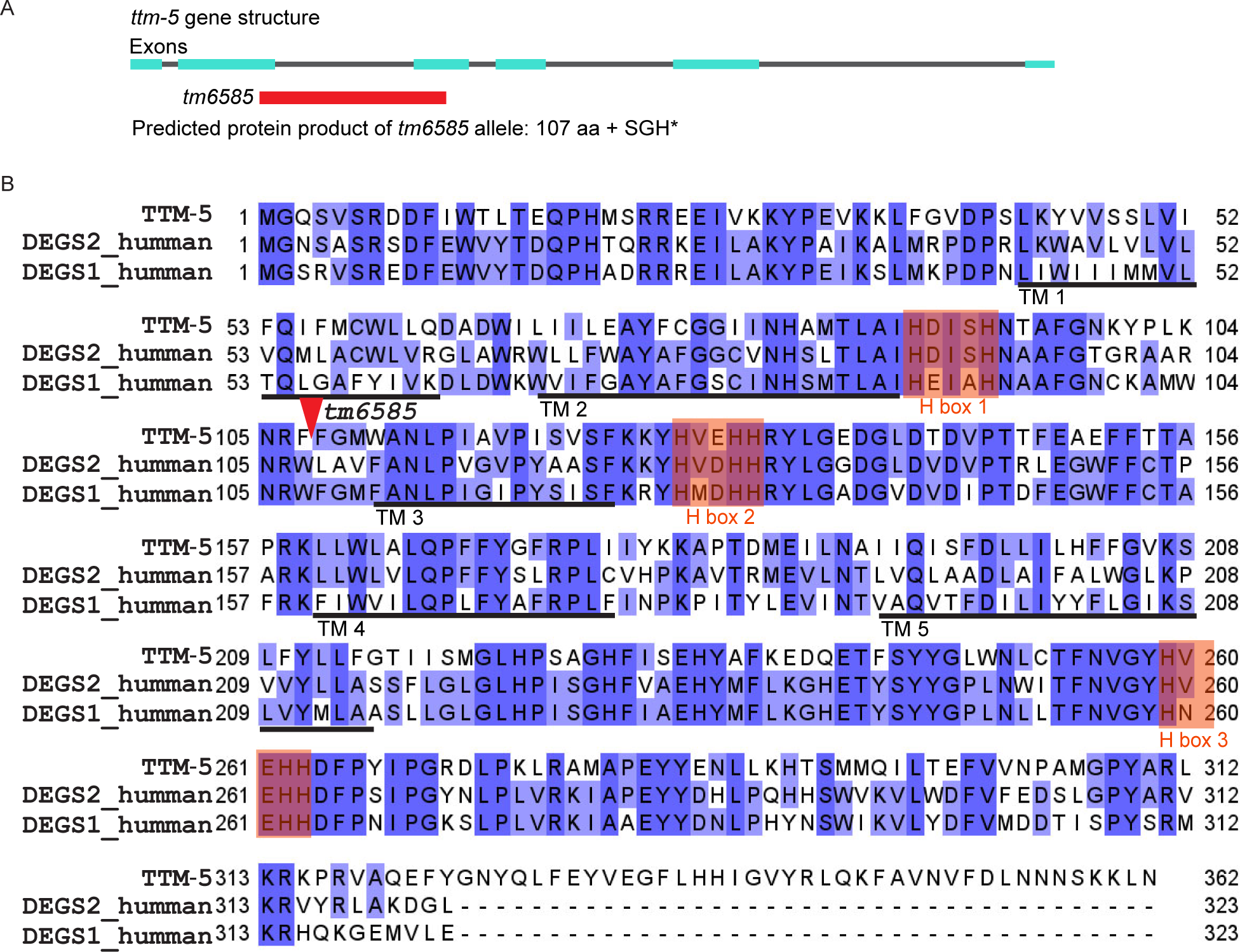
*ttm-5 (tm6585)* lesion analysis, related to **Figure 4**. (A) The deletion allele *tm6585* removes sequences between exon 2 and exon 3, resulting in a truncated protein that retains 107 amino acids of the TTM-5 protein (indicated by red arrow in panel B), followed by three frame shift amino acids. (B) Alignment of TTM-5 with its human orthologs DEGS1/2. TTM-5 and DEGSs are predicted to have five transmembrane domains. The allele *tm6585* is a null allele that eliminates two H boxes, which are histidine-rich motifs crucial for desaturase activity [S5].

